# Measuring stepwise binding of a thermally fluctuating particle to a cell membrane without labeling

**DOI:** 10.1101/763680

**Authors:** A. Rohrbach, T. Meyer, H. Kress

## Abstract

Thermal motions enable a particle to probe the optimal interaction state when binding to a cell membrane. However, especially on the scale of microseconds and nanometers, position and orientation fluctuations are difficult to observe with common measurement technologies. Here we show that it is possible to detect single binding events of IgG-coated polystyrene beads, which are held in an optical trap nearby the cell membrane of a macrophage. Changes in the spatial and temporal thermal fluctuations of the particle were measured interferometrically and no fluorophore labelling was required. We demonstrate both by Brownian dynamic simulations and by experiments that sequential step-wise increases in the force constant of the bond between a bead and a cell of typically 20 pN / µm are clearly detectable. In addition, this technique provides estimates about binding rates and diffusion constants of membrane receptors. The simple approach of thermal noise tracking points out new strategies in understanding interactions between cells and particles, which are relevant for a large variety of processes including phagocytosis, drug delivery or the effects of small microplastics and particulates on cells.

**SIGNIFICANCE:** Interactions of cells with nearby particles, e.g. bacteria, viruses or synthetic material, is a very fundamental and complex process, often deciding about the cellular fate. The investigation of binding processes between particle and cell is typically investigated by fluorescence techniques, where fluorophores often hinder the molecular interaction of specific binding partners. Therefore, label-free detection or imaging techniques are essential, which are hardly available especially for live cell investigations. Molecular binding is based on thermal position and orientation fluctuations of the binding partners to find the best interaction state. Here, we present a label-free measurement technique that allows us to detect multiple stepwise binding events of molecules on an optically trapped particle close to the cell membrane.

## INTRODUCTION

The endocytic uptake of particles into cells is controlled by multiple biochemical and biophysical mechanisms. The most prominent cellular process, phagocytosis [1-3], comprises the engulfment, internalization and intracellular transport of a particle, requiring energy for the deformation of the cell membrane, the reorganization of the actin network [4] and for phagosome transport by molecular motors [5, 6]. In many cases endocytosis is receptor-mediated, which means that ligands on particles, typically bacteria and viruses, trigger the recruitment of receptors in the cell membrane, such as the Fc receptor for immunoglobulins (Ig) or receptors of the complement system. The role of the spatial distributions of receptors, their diffusivity and their kinetics of ligand binding are of superior interest to understand their dynamic function not only during endocytosis [7], but also during other fundamental cellular processes such as the formation of immunological synapses during antigen presentation [8] or the formation of focal adhesions [9].

However, the experimental investigation of such processes is difficult since they take place on very small length and time scales. In addition, established optical techniques such immunocytochemical staining, Förster resonance energy transfer (FRET) and fluorescence recovery after photobleaching (FRAP) provide usually strong signals, but often may influence the receptor ligand binding because of attachment of fluorophores or because they are based on genetic modification of membrane proteins [10].

Against this background, various label-free optical detection methods have been developed or applied, based on assay approaches using changes in local refractive indices within the membrane [11-14] or AFM-based single molecule studies [15] Whereas many mechanistic insights into working and design principles of biomolecules are averaged out by bulk assay methods, single molecule approaches provide more detailed information on both spatial and temporal scales. A large amount of information can be extracted when the thermal fluctuations between the binding partners are not suppressed, but are measured on broad bandwidths, with e.g. Photonic Force Microscopes (PFM) [16], and are analyzed with correlative or multi-spectral methods ([17, 18][19]).

A PFM [20] is an optical tweezers-based apparatus being able to bring binding partners in close vicinity of each other with help of an steerable optical trap and to track the thermal motions of one or both binding partners in parallel. Using back-focal plane (BFP) interferometric tracking [21], the motions of a single bead [22] or bacterium [23] or several particles [24] can be recorded in three dimensions with nanometer precision and at rates of up to 2 MHz. Studies using PFM have investigated individual streptavidin–biotin complexes on functionalized surfaces revealing sequential bond formation [25], force-spectral bond ruptures [26] or intermediate states during membrane fusion [27].

In our study, we were able to expand this approach to investigate the successive binding of individual bonds of IgG ligand coated optically trapped beads and Fc-γ receptors in the plasma membrane of a living mouse macrophage.

### Experimental and mechanistic principle

A 1 µm sized polystyrene (PS) bead coated with immunoglobulin G is approached by an optical trap to the periphery of an adherent J774 mouse macrophage. Care was taken to choose flat areas of the cell surface without any visible filopodia, here imaged by differential interference contrast (DIC) microscopy (see Figure 1). In a first stage, the bead diffuses inside the stationary optical trap without being affected by the presence of the cell as outlined by the sketch of Figure 1C. After some seconds, the bead’s motion is significantly changed in space and time because of subsequent binding of possibly IgG ligands to the Fc-γ receptors in the plasma membrane. The change in the position fluctuations can hardly be detected with conventional contrast enhancing microscopy methods (such as DIC), but can be tracked by back focal plane interferometry very precisely in three dimensions. The molecular scale fluctuation analysis of the temporal and spatial change of the bead center positions, which were measured by PFM on the one hand and simulated through Brownian dynamics (BD) on the other hand, define the underlying concept of the study. The mechanistic model as depicted in Figure 1E,F describes the relevant constituents: the optically trapped bead, the surface ligands, the membrane receptors and thermal driving forces. The temporal evolution of these counteracting forces can be described by a mathematical force equation, which represents the basis for our data analysis and the numerical solution of this force equation via BD simulations.

**Figure 1:**
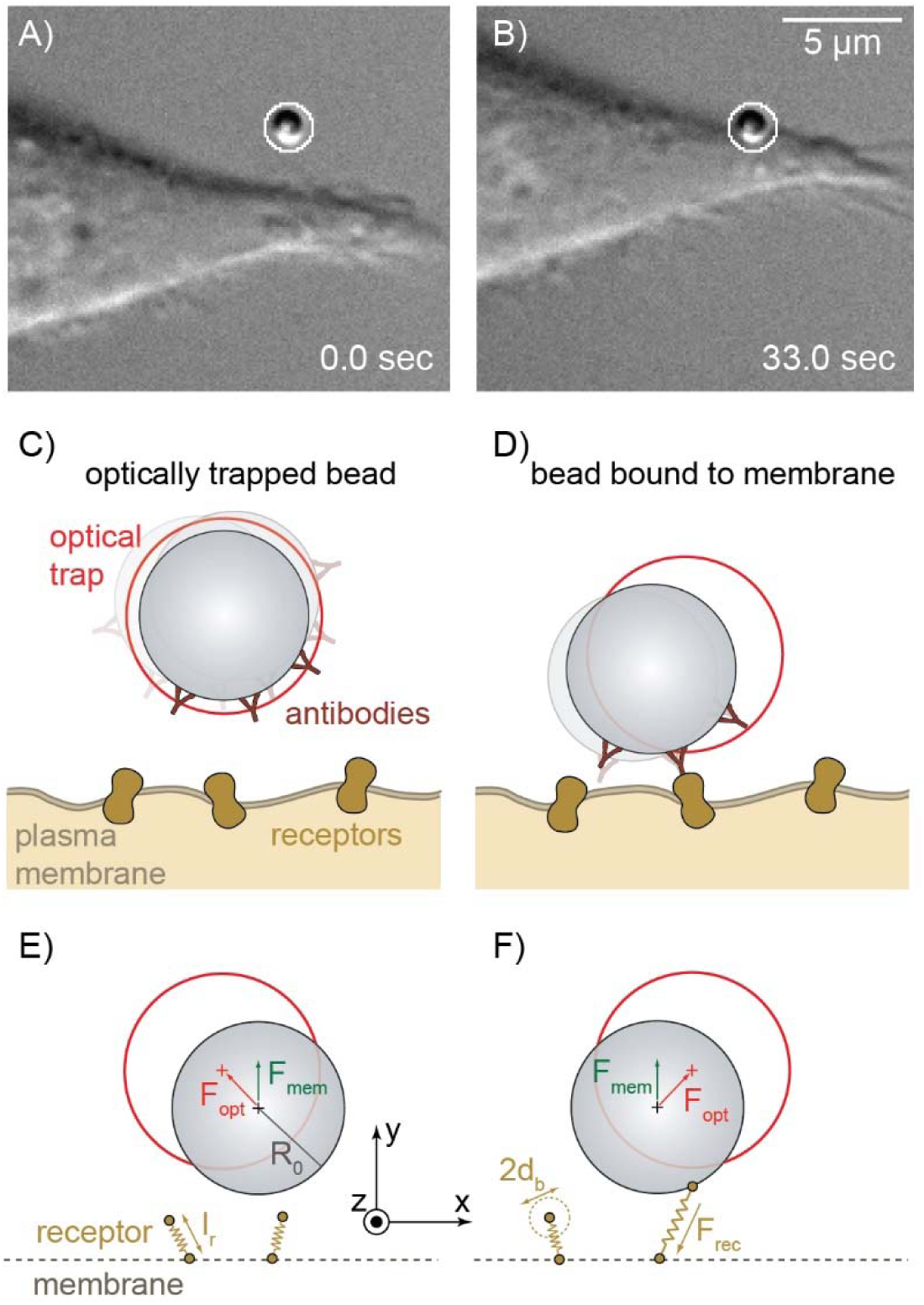
Measurement of the thermal motion of a microparticle during binding to a cell membrane. (A), (B) DIC images of an optically trapped IgG-coated microparticle before (a) and after (b) binding to the membrane of a macrophage. (C, D) Sketch (not to scale) of the optically trapped antibody-coated particle before (c) and after (d) binding to receptors in the cell membrane. (E,F) Schematic of the Brownian dynamics simulation of the binding process showing the particle in the optical trap before (E) and after (F) binding to receptors in the cell membrane.

## MATERIALS AND METHODS

### Cell culture and handling

The cells used for the experiments were J774A.1 murine mouse macrophages (17). Cells were cultivated as described previously (Kress et al., 2007). One day before the experiments, the cells were seeded on coverslips.

To assure the access for the optical tweezers an appropriate confluency is important. A cell coverage between 10% and 30% of the coverslips at the day of the experiments turned out to be reasonable. In order to provide physiological conditions during the experiment the cells were kept heated at 37°C by a custom-built heating system.

### Optical trapping and interferometric tracking

The experiments were performed on a self-developed Photonic Force Microscope (PFM). The microscope was equipped with a 3D piezo stage and is extended by the standard units for optical trapping and tracking (18), such as a 1064 nm laser (Crysta-Laser).

The binding process was studied by tracking the 3D position of the bead using back focal plane (BFP) interferometry at frequencies of 10 to 100 kHz. Before every experiment, the optical trap and the QPD detection system were calibrated with the methods described in (18–20). Scattering of the trapping laser light at the cell affects the mean value of the position signal, but not the fluctuation width of the position signal [18] and therefore need not to be considered in our analysis.

### Mathematical description of successive particle binding to the cell

In this section, we translate our mechanistic picture sketched in Figure 1 into a mathematical description of the particle motion based on differential equations (DEs), which consider the most important forces influencing the particle. On the one hand, the DEs will be solved analytically to give a basis for the correlation analysis methods. On the other hand, the DEs will be solved in more detail numerically by Brownian dynamic simulations described in the second part of this section.

The bead’s stochastic motion in the vicinity of a cell surface can be described by an overdamped Langevin equation

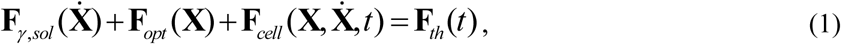

which we use to analyze and identify the cell binding characteristics. Here, the state vector **X**(*t*) = (x,y,z,α,β) describes both the center position **r**(*t*) = (x,y,z) and the orientation **θ** = (α,β) of the bead relative to the trap center at a given time *t*. **F**_*γ*, *sol*_ is the frictional force of the bead in the surrounding fluid, **F**_*opt*_ the optical force and **F**_*cell*_ the force, which the cell with membrane receptors exerts on the bead. The system is driven by a random thermal force **F**_*th*_, which accounts for the Brownian motion of the particle.

The optical force is approximated to be linear: 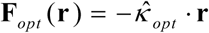, where 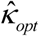 is a diagonal matrix containing the direction-dependent stiffnesses of the optical trap. The frictional force 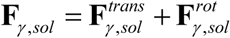 can be expressed by 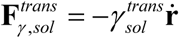 and 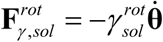, where 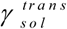 and 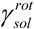 are the translational and rotational friction coefficients of the bead in the surrounding fluid solution.

For the sake of readability and to concentrate on the relevant message of this paper, the analytical model used for the receptor number and the autocorrelation analysis neglects the bead rotation, which is however considered in the Brownian dynamics simulations.

In first-order approximation, the binding of the bead to the membrane receptors is assumed to be harmonic. A single ligand-receptor bond is described by a time-varying harmonic potential and thus an elastic force **F**_*κ,rec*_ (**r**,*t*)=−*κ*_*rec*_ ·(**r**(*t*)+**R**_0_(*t*)), where *κ*_*rec*_ denotes a time-independent isotropic stiffness of the receptor at position **r**_*rec*_(*t*) =**r**(*t*)+**R**_0_(*t*), **R**_0_ is vector from the bead center to the receptor position. In addition to the elastic binding force, the ligand-receptor bond exerts a frictional force onto the bead, expressed by the linearized force 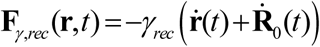, where *γ*_*rec*_ is an isotropic molecular friction coefficient influencing the temporal fluctuations of the bound bead. Hence, the receptor changes its position by diffusion and through the elastic force of its bond with the bead. For an increasing number *N*(*t*) of receptors (index *n*) that bind over time to the bead surface and successively change the fluctuation behavior of the bead, the linearized Langevin equation (1) for the bead position can be expressed by

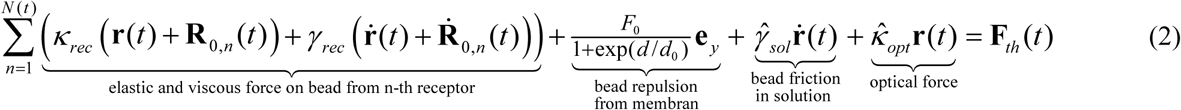

To avoid an unrealisitic transition of the bead through the membrane, a sigmoidally smoothed hard-core repulsion force *F*_*mem*_·**e**_**y**_ of the bead at the membrane was introduced. This membrane force is 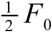, when the distance d to the bead surface (≈ d+R_0_ to the bead center) reaches the membrane deformation length d_0_ ≈ 20 nm, which is on the order of about 2-3 times the membrane thickness. At length d_0_, the slope of F_mem_ is 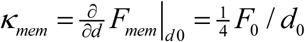.

Upon binding the center position of the bead with radius R_0_ mainly shifts in y direction by the height h(t), which is the direction radial to the cell surface. Therefore, the state vector is approximated to **r**(*t*) = (0, *R*_0_ + *h*(*t*),0,0,0) to estimate the number of receptors.

### Number of receptors binding to the particle

Here, we describe a particle binding behaviour prior to an active formation of the phagocytic cup. We use a simple, empirical model to describe the increase in binding area *A*_*cap*_(*t*) through a passive, spherical cap shaped indentation and the concentration increase *c*(*t*) of the receptors (transmembrane proteins such as FCγ receptors) by fusion and binding to the spherical surface of the bead. In our simple model, both *A*_*cap*_ (*t*) and *c*(*t*) increase independently of each other in time.

In eq. (2), the number *N* (*t*) of membrane receptors bound to the bead

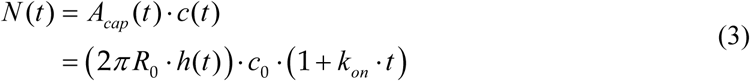

is assumed to increase with time both due to an increase of the contact area of the spherical cap, *A*_*cap*_ (*t*) = 2*π R*_0_ · *h* (*t*) with indentation height h, and the receptor concentration *c*(*t*).

The temporal increase of the indentation height *h*(*t*) (and thereby the spherical cap area *A*_*cap*_ (*t*)) is modeled by an exponential *h*(*t*) = *h*_max_ (1−exp(*-t* / *t*_*0*_)), which approaches a maximum indentation height *h*_*max*_. This value is much smaller than the bead radius, i.e. *h*_*max*_ < *R*_0_, and is reached when the adhesion energy between bead surface and membrane equals the deformation energy of both membrane and actin cortex. Our two-parameter approach is similar to a published power-law model [28, 29], which was used for modelling the formation of a phagocytic cup.

We further assume an initial receptor concentration c_0_ at time t = 0 and, as a first-order approximation, a linear increase in receptor concentration with time *c*_0_ ·*k*_*on*_ ·*t* through free diffusion. *c*(*t*) can vary with each cell. The receptors are assumed to bind to the bead with a rate *k*_*on*_, which is in general limited by diffusion and by fluctuations. Furthermore, we assume that receptor unbinding is negligible. On the timescale of seconds, we assume that the contacting process is only diffusion-limited and molecular binding occurs instantaneously, such that the binding rate 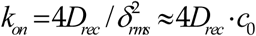 depends on the diffusion constant *D*_rec_ of the receptors in the membrane and the root-mean-square distance 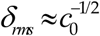 between the receptors. In this approximation, the time-variant number of receptors in eq. (3) can be refined to

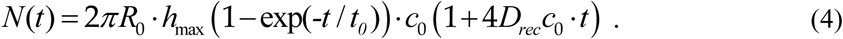

Here, 4*D*_*rec*_*t* is the mean square displacement (MSD) for two-dimensional free receptor diffusion with *D*_*rec*_ = *k*_*B*_*T*/*γ*_*rec*_ defined by the receptor friction coefficient *γ*_*rec*_ introduced above.

Solutions of eq. (4) for various parameters will be compared to both experimental results and BD simulations and are presented in Figure 6.

### Increase in binding stiffness and friction

A suitable and common method to determine forces constants κ and friction coefficients γ from fluctuation data of beads is correlation analysis [30]. The bead experiences the harmonic potential of the optical trap and the sum of the harmonic potentials of each bound receptor, of which the number increases during the measurement time *t*_*m*_. The resulting potential is again harmonic and can be described by the total stiffness or binding strength κ_tot_(*t*_*m*_), which is the sum of the trapping stiffness κ_opt_ and the increasing sum of all receptor stiffnesses N(*t*_*m*_)·κ_rec._ The total friction factor γ_tot_(*t*_*m*_) is the sum of friction factors experienced by the bead in the solution γ_sol_ and in contact with the receptors, N(*t*_*m*_)·γ_rec_.

Neglecting the directional dependency for a moment, the increase in binding strength and friction can be summarized as follows

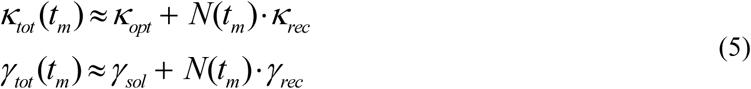

Both values can be measured for a small number N of receptors through autocorrelation analysis.

### Autocorrelation analysis

Assuming a linear force and friction acting on the particle in thermal equilibrium (see eq. (2)), within a short time-window the autocorrelation function 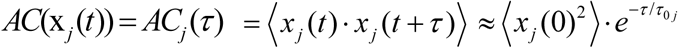 for the particle’s position in direction j = x,y,z is given by:

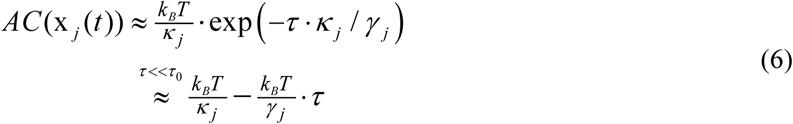

Applying a two-parameter exponential fit to the second line of eq. (6), each 3 parameters *κ*_*j*_ for stiffness and friction *γ*_*j*_ can be extracted, by determining the AC time *τ*_*0j*_ = *γ*_*j*_/*κ*_*j*_. For time delays *τ* much shorter than *τ*_*0j*_, the autocorrelation can be well approximated to be linear in τ. The static part (τ = 0) of the AC yields the stiffness *κ*. The dynamical information extracted from the slope 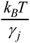 of the AC for short time delays τ yields the friction coefficient γ. The two parameters described in eq. (5) can be analyzed within a time-window Δt > *τ*_*0j*_ for each time point *t*_*m*_ as follows:

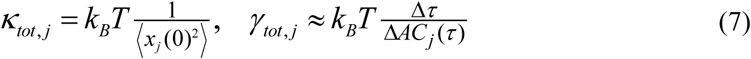

The stiffness can be obtained both from equilibrium information over the fluctuation width σ_*j*_ = ⟨ *x*_*j*_ (0)^2^ ⟩^1/2^ or through the exponential decay time τ_0_ of the position autocorrelation function.

### Brownian Dynamics Simulation

We used Brownian dynamics simulations to investigate the binding of a microparticle to the membrane of a cell as depicted in Figure 2A. Particles immersed in aqueous solution experience fluctuating thermal forces due to collisions with the water molecules. The autocorrelation time *τ*_*w*_ of these fluctuations is on the order of the average intermolecular separation divided by the mean molecular velocity, which yields about 10^−13^ s at room temperature [31]. On time scales, which are large compared to *τ*_*w*_ the motion of the centre-of-mass coordinate *x*_*j*_ of a particle is described by the Langevin equation in the low Reynolds number limit. The translational and rotational Brownian motion of the particle is

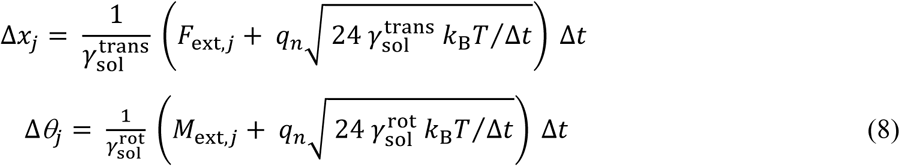

where in the upper equation *F*_*ext*_ summarizes all additional deterministic external forces which act on the particle besides the translational viscous drag forces and the random thermal forces. In the lower equation, *M*_th_ represents the thermal torques acting on the bead and *M*^ext^ summarizes all additional deterministic external torques besides the rotational viscous drag torques. The amplitude of the thermal force is described by the fluctuation-dissipation theorem 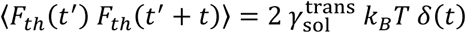 where *δ* is the Dirac delta distribution (thermal torque M_th_ analogous). To simulate the Brownian motion of a particle, the thermal forces (and torques) can be calculated by using a random number generator. For evenly distributed random values *q*_*nn*_ = [-0.5 … 0.5] the maximal amplitude of the thermal force is [32] 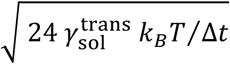, where *Δt* denotes the temporal resolution of the simulation.

**Figure 2:**
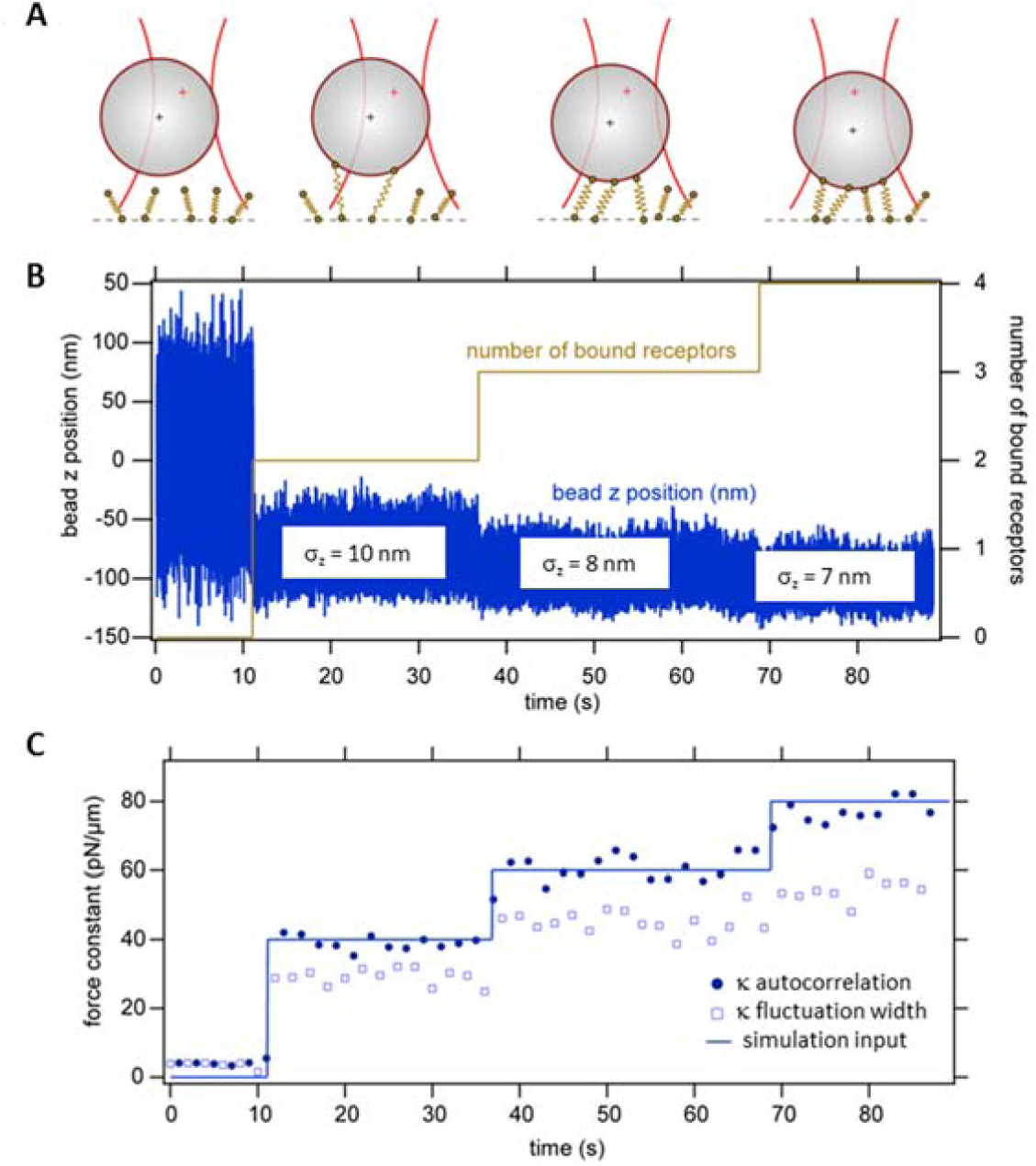
Brownian dynamics simulation of sequential receptor binding. An optically trapped bead is moved towards a cell and binds sequentially to membrane receptors. (A) Sketch of the sequential binding process. (B) Bead position trace in z-direction. (C) Total force constant *κ*_*tot*_ determined by the autocorrelation time and the fluctution width.

## RESULTS

### Simulations

The 1 µm sized particle was simulated to diffuse in the 3D harmonic potential of an optical trap with typical trap stiffnesses *κ*_*x*_ = *κ*_*y*_ = 20 pN/µm and *κ*_*z*_ = 4 pN/µm [33]. Membrane receptors were embedded in the membrane at an effective density ranging around *c*_*0*_ = 4/µm^2^, assuming 100% binding probability. Since we do not consider the probability of receptor binding, which is significantly less than 100%, we use an effective receptor density, which is lower than the densities reported in the literature [34]. The receptors were allowed to diffuse freely in the plane of the membrane with diffusion coefficients ranging between *D*_*rec*_ =0.03 µm^2^/s and *D*_*rec*_ = 0.07 µm^2^/s. The mechanics of a receptor was simulated in a first order approximation as a linear spring with a spring constant of *κ*_*rec*_ = 20 pN/µm as described in eq. (2). The orientation of the receptor was not restricted, but it was held in the membrane by a harmonic potential in the z-direction (perpendicular to the plane of the membrane) with a force constant *κ*_*mr*_ = 20 pN/µm reflecting the embedding of the receptor in the membrane.

During the time course of the simulations, the particle was first fluctuating in the optical trap without any contact to the cell membrane. The trap was then moved slowly towards the cell until the bead was bound to the membrane. Binding to the membrane through a receptor was initiated when the surface of the bead was sufficiently close to the top part of the receptor (binding distance *d*_*b*_ = 20 nm). After one or more receptors bonds were formed, each receptor exerted a force and a torque onto the particle as depicted in Figure 2A.

Figure 2B shows an examplary simulated trajectory of a particle that binds over the time course of several tens of seconds sequentially at first to two receptors simultaneously, then to a third receptor and subsequently to a fourth receptor. All simulations were performed in three dimensions, but only the z-direction is shown here for simplicity.

It can be seen in the particle trajectory (Figure 2B) that the width of the fluctuations decreases with the first and second binding event. A quantitative analysis of the fluctuation width *σ* is shown in Figure 2C. In time intervals of 2 seconds, the standard deviation *σ* of the particle position fluctuations was calculated with a time window of Δt = 2s and the corresponding total force constant of the binding *κ* was calculated by 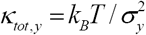 according to eq.(7). This force constant (“binding strength”) is in a first approximation the sum of all the bound receptor stiffnesses. Since the optical trap and the membrane receptors form a parallel arrangement of damped springs, the optical trap stiffness provides a constant offset for *κ* and can be easily. The expected total force constant (labeled “total receptor *κ*”) is the sum of the bound receptor force constants, i.e. 40 pN/µm for *N*(*t*_*m*_) = 2 bound receptors, 60 pN/µm for *N*(*t*_*m*_) = 3 and 80 pN/µm for *N*(*t*_*m*_) = 4bound receptors, at times *t*_*m*_ = 11s, *t*_*m*_ = 38s and *t*_*m*_ = 70s, respectively. This corresponds to fluctuation widths σ_z_ of about 10nm, 8nm and 7nm, which can be distinguished. However, it can be seen that the σ_z_ analysis underestimates the parameter *κ*. One reason for this underestimation is that the receptors are assumed to be held in the membrane by a harmonic potential in the y-direction, i.e. perpendicular to the membrane. The resulting small diffusion of the receptors in z-direction increases *σ* and therefore decreases *κ*.

Instead of determining the total binding strength *κ* via the fluctuation width *σ*, it can also be calculated via the autocorrelation time *τ* of the fluctuations according to second line of eq. (6). The result of this type of analysis is also shown in Figure 2C (filled circles). If the characteristic time of the receptor motion in the membrane potential separates sufficiently well from the characteristic time of the bead motion, this method for deriving *κ* gives more precise values. This can be seen by the good agreement between *κ* (autocorrelation method) and the expected value.

The results of the BD simulation presented in Figure 2 demonstrate that it is in principle possible to extract single molecular binding events from 1µm sized beads fluctuating in position and orientation. The result is not obvious since our thermal noise analysis bases only on the fluctuations of the bead center positions, disregarding the orientation fluctuations and the torques induced by the receptors binding to the bead surface in a distance *R*_0_ = 500nm. The result is further surprising since the molecular springs, representing the receptors, also act in oblique direction, but only the vertical components alongside directions are analyzed.

To enable further comparisons with experimental results (see next section), we ran the BD simulations repetitively over 12-20 seconds, to simulate the number *N*(*t*) of receptors binding with time for the above mentioned membrane concentrations c_0_ and receptor diffusion coefficients D_rec_. Figure 3B displays 30 receptor binding traces *N*(*t*_*m*_), which increase first nonlinearly, then linearly and stepwise with time. The colored traces indicate the average receptor numbers 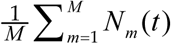, where individual steps are no longer visible and which are summarized in Figure 3A. Here, the standard deviation is shown exemplarily only for the red trace.

**Figure 3:**
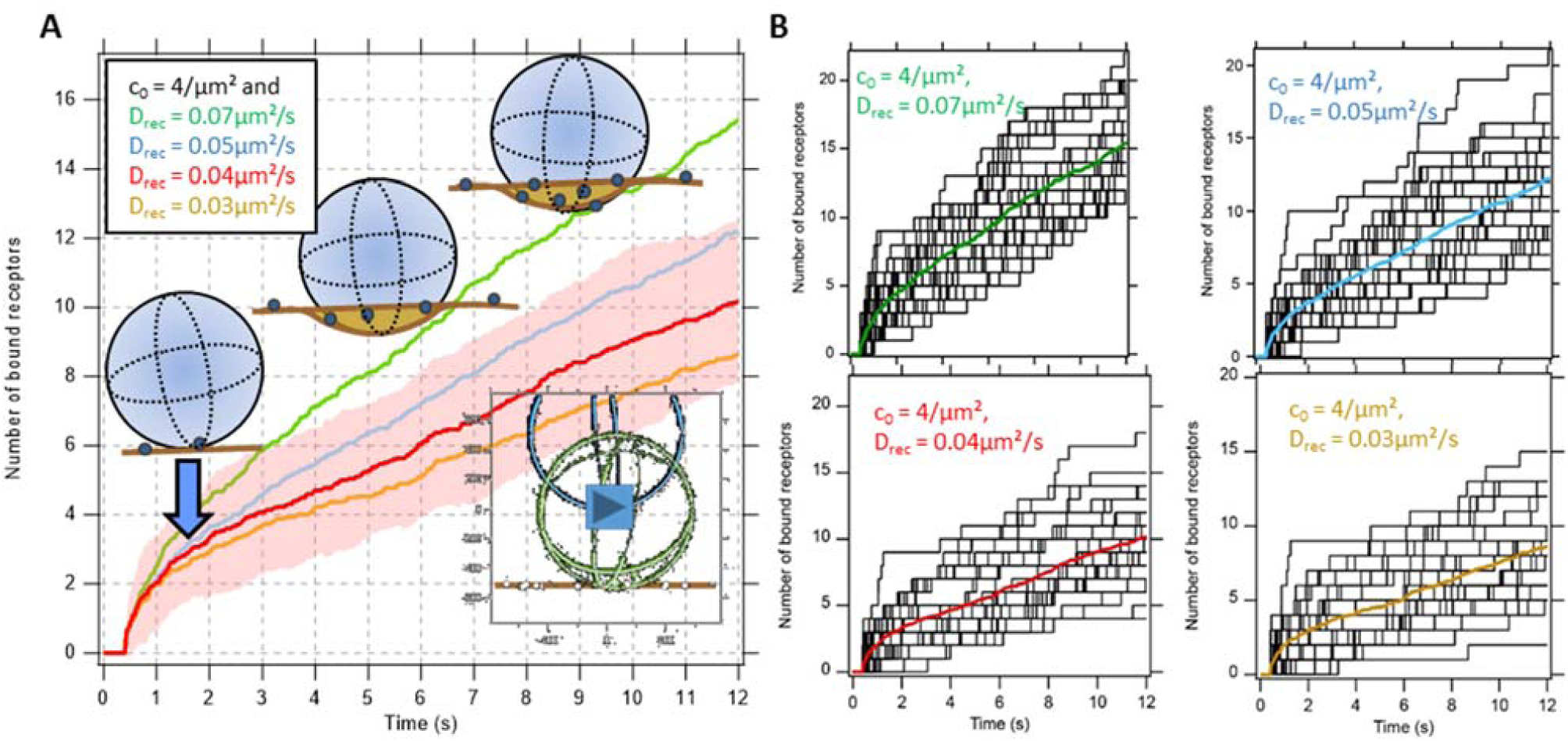
Brownian dynamics simulations of receptor binding as a function of time t_m_. Model parameters were K_rec_= 20 pN/µm for single receptor binding strength and initial concentration c_0_ = 4/µm^2^, D_rec_ = 0.03…0.07 µm^2^/s. (A) Mean number of bound receptors during subsequent stages of bead indentation and receptor diffusion. See supplementary movie. (B) Stepwise binding of single receptors on bead and analyzed from BD simulations (black tracks) and averaged trace in color.

### Experiments

In a series of experiments, we measured the time course of the binding of IgG-coated microparticles to the membrane of macrophages. We trapped IgG-coated polystyrene spheres and moved them in close vicinity of about 100 nm to a visibly flat membrane of a macrophage (see Figure 1). The thermal fluctuations of the particle’s position and orientation allowed the IgG ligands and the Fc-receptors in the membrane to find a preferable binding condition. We measured the thermal motion of the bead center positions before and during the binding event by BFP interferometry with a spatial precision of several nanometers and a temporal resolution between 10 µs and 100 µs. A representative data set of the thermal motion of a particle during binding to a cell membrane is shown in Figure 4A, which typically does not allow to detect significant changes in the fluctuation behavior by pure eye observation.

**Figure 4:**
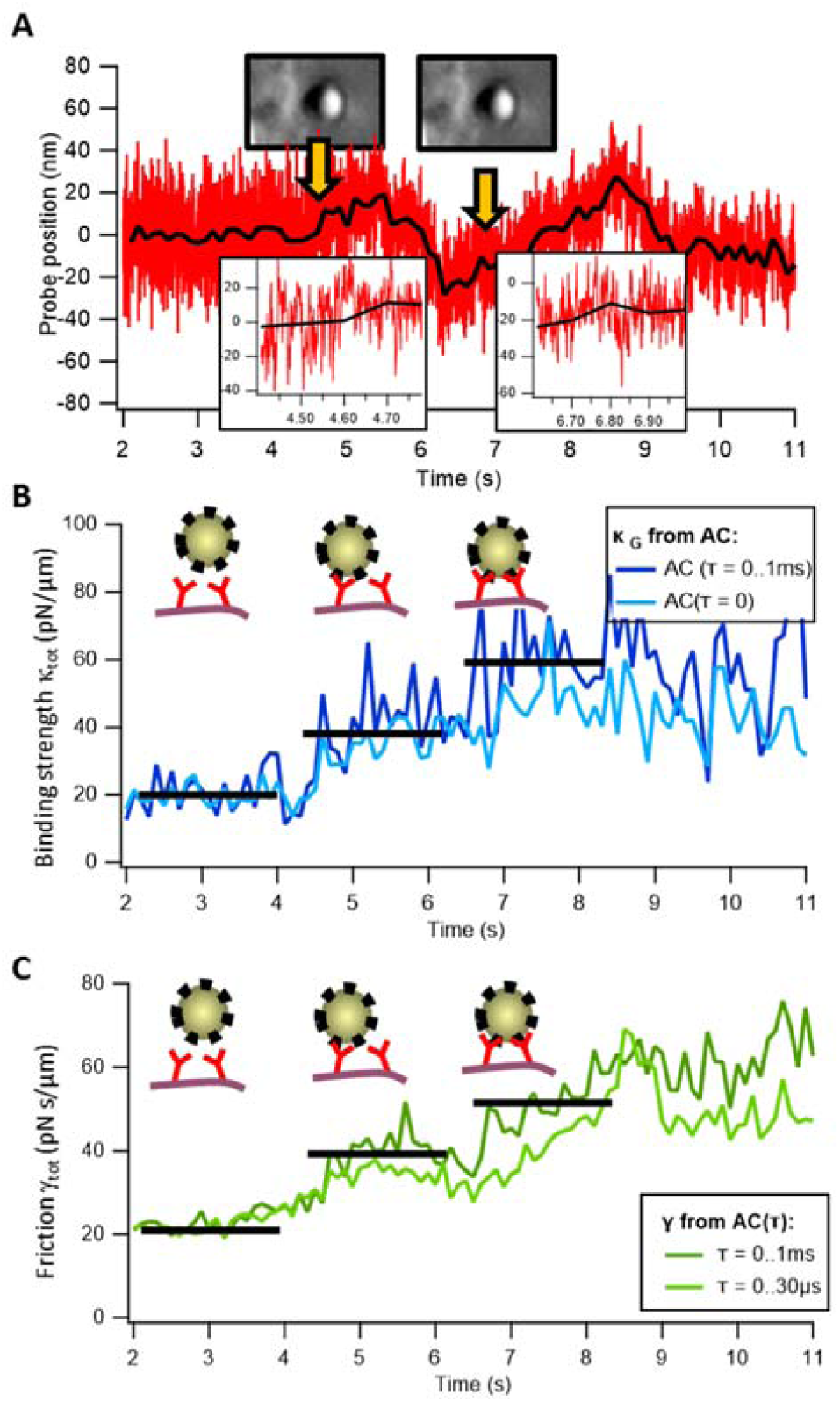
Binding dynamics between IgG-coated microparticles and cells. (A) Thermal motion of an optically trapped IgG-coated particle that binds stepwise to the membrane of a macrophage (see orange arrows and insets), which become visible in the plots below. (B) Autocorrelation analysis of the particle position fluctuations allows determining the binding force constant κ_tot_(t). The binding force constant increases stepwise as the particle binds to the cell. A second abrupt increase a few seconds after the initial binding to the cell indicates a further bond formation between particle and cell membrane. (C) Stepwise change of the viscous drag γ_tot_(t) experienced by the particle upon binding.

However, the analysis of the autocorrelation of the particle’s position fluctuations allows the determination of the total binding force constant κ_tot_(t_m_) and the friction constant γ_tot_(t_m_) as a function of measurement time t_m_ (see eq. (5)). This binding force constant (binding strength) is the sum of the optical trap stiffness κ_opt_ and the force constants of the receptors κ_rec_(t_m_), the friction constant reflects the damping of the bead at the membrane and the connection to the molecular spring with γ_rec_(t_m_), which both describe the binding to the cell in a two parameter mechanistic model as introduced in the theory section.

Figure 4B,C show the temporal course of κ_tot_(t_m_) and γ_tot_(t_m_), which fluctuate depending on the strength of averaging and the chosen window size for each measurement point t_m_. However, it can be seen in Figure 4B that the binding force constant fluctuates around a plateau value, which increases abruptly as the particle binds to the cell (to a value at around 40 pN/µm). About two seconds later, a second abrupt increase (the third plateau value at around 60 pN/µm) is visible, indicating an additional bond formation between the particle and the cell. Corresponding increases at related time points are visible in the time-course of the friction constant in Figure 4C. Both binding events are marked by orange arrows in Figure 4A.

Based on the principles described in eqs. (6) and (7), the varying decay of the position autocorrelation reflects the viscous drag γ experienced by the particle before binding, after the first binding and after the second binding to the membrane. Both the binding force constant and the friction constant reveal a stepwise increase, which have been analyzed with the two different types of analysis. As demonstrated by the results of the BD simulations, the analysis of the whole correlation decay provides a reliable set of parameters.

A later series of experiments at higher sampling rate (at 1 MHz) allowed to analyze both the stepwise increase in binding strength and friction as demonstrated by the exemplary results of Figure 4.

In an earlier series of experiments at a lower sampling rate (10 - 50 kHz), we performed binding experiments in which optically trapped IgG-coated particles of the same size were placed in close vicinity to the membrane of J774 macrophages. As shown in Figure 5A, we determined the initial stepwise increase of the binding force constant in experiments, where this was visible. In total, we identified 45 stepwise increases in the binding strength. Figure 5A displays the temporal course of the binding stiffness κ_tot_(t_m_) (using two different smoothing values) revealing two discrete steps with a height of about 35 pN/µm and 50 pN/µm. Before the time point t = 34 s, the particle was optically trapped with a force constant of about 50 pN/µm. Analysis of all 45 binding events by fitting the multi-Gauss function 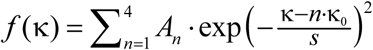 results in a distribution of force constant jumps *n* · κ_0_ = *n* ·(19 ±1) pN/µm as shown in Figure 5B, with s = (11±1) pN/µm and **A** = (13,2,1,1) ± 1. Again, we associate thes jumps in force constant with the binding of the particle to an individual Fcγ-receptor.

**Figure 5:**
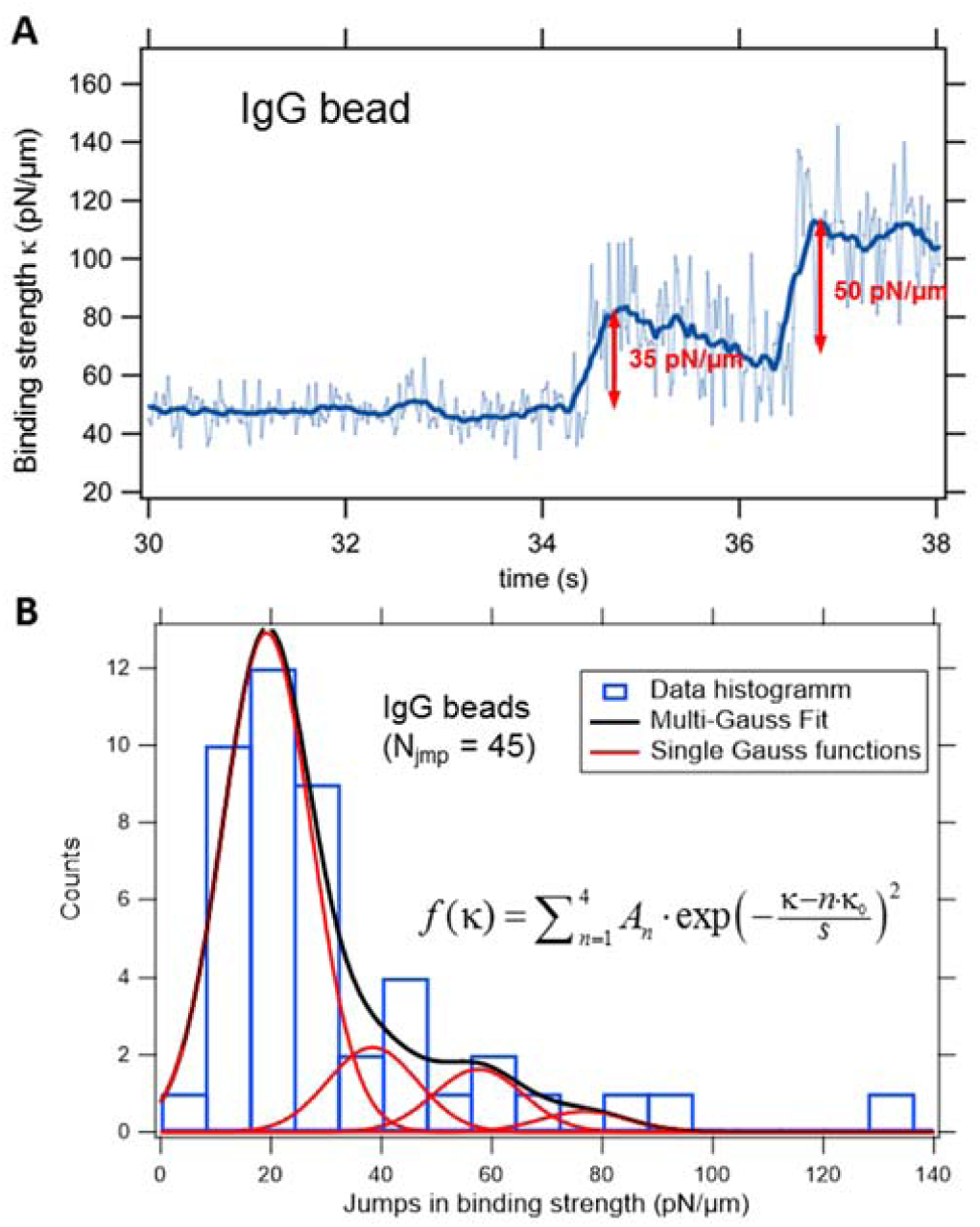
Stepwise increase of the binding strength κ_tot_(t) during binding of IgG-coated particles to J774 macrophages. (A) Temporal course of binding force constant with two distinct steps. (B) Histogram of the observed stepwise increases of the binding force constants. The total number of measured jumps in force constant was N_jmp_ = 45. The histogram from the multi-Gaussian fit has peaks at n·κ_0_ = n·19 pN/µm.

In addition to the stepwise increase of binding strength because of individual bond formation events, it is possible to determine the mean increase in binding strength between particles and cells as a function of time, by averaging all measured time-courses of the binding strength constants. Figure 6A shows the mean of the binding strength, which increases with time during the first ten seconds after the initial binding event. Whereas the trajectories of single experiments fluctuate strongly, the average of 45 experiments shows a nearly linear increase with time, where the slope slightly depends on the averaging strength. For four different smoothing parameters, corresponding to time windows Δt = 25ms…200ms, the slope varies by not more than about ± 20%. The association of an increase in binding strength of κ_rec_ = 20 pN/µm with each additional receptor leads to an average binding of N = 11 receptors during the first 10 seconds of particle cell interaction (at Δt = 25ms). Using a broad smoothing window of Δt = 200ms, the analysis provides that N = 7 receptors bind in average within 10s (Figure 6, right ordinate). For a reported receptor density of c_0_ = 10…100 / µm^2^ with limited binding probability [34], we approximated c_0_ ≈ 4 / µm^2^ assuming 100% binding probability in the BD simulation.. In the analytical model, the cap area is A_cap_(t_m_=1s) = 0.1µm^2^ with receptor density c_0_ ≈ 0.4 / 0.1µm^2^ in the first second and increases to N = 10 receptors after 10 seconds with a contact area A_cap_(10s) = 0.16 µm^2^, such that c(10s) ≈ 63 / µm^2^.

**Figure 6:**
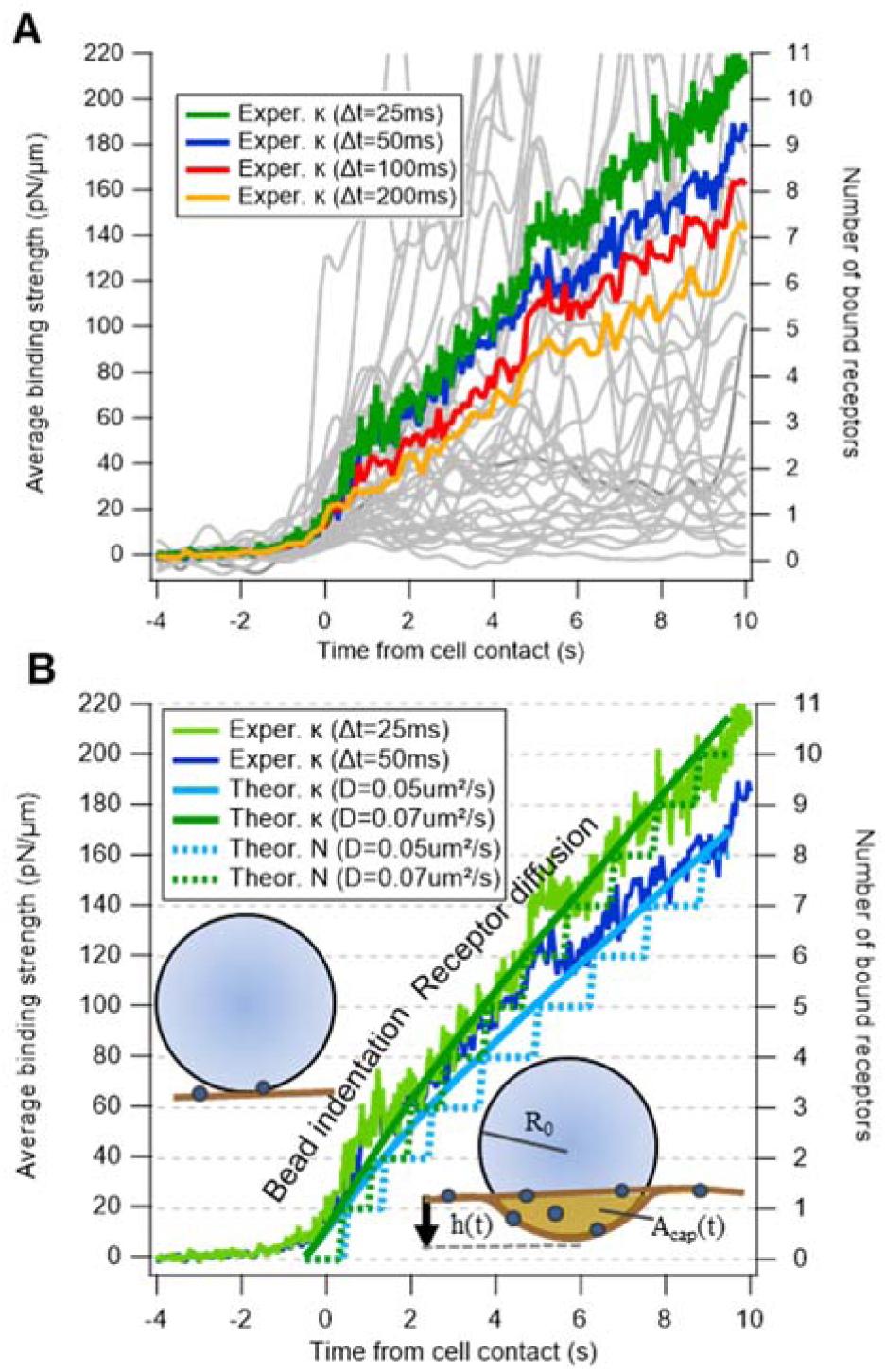
Increase in binding strength between IgG-coated particles and J774 macrophages after particle-cell contact. (A) Measured binding strength traces κ_tot_(t)-κ_opt_ for 45 different contacts (light gray) smoothed over Δt=100ms time window. Averaged binding strength smoothed over time windows Δt = 25ms…200ms. (B) Two averaged binding strengths relative to traces κ_tot_(t)-κ_opt_ obtained by the theoretical model. An assumed single receptor binding strength of κ_rec_ = 20 pN/µm leads to average binding of 10-11 receptors within 10 seconds (right ordinate). Model parameters: h_max_ = 80nm, c_0_ = 4/µm^2^, D_rec_ = 0.05 µm^2^/s (0.07 µm^2^/s), R_0_ = 0.5µm, t_0_ = 1s.

Using the initial concentration c_0_ ≈ 4 / µm^2^, coincidence with the binding model described by eq. (4) is achieved for a receptor diffusion constant of D_rec_ = 0.05 .. 0.07 µm^2^/s, which is at the lower limit in a list of experimentally obtained receptor diffusion constants[35], mainly measured by FRAP (fluorescence recovery after photobleaching). A comparison between results obtained from experiments and theory are shown in Figure 6B for each two sets of curves, which all show a stronger increase directly after contact t_m_ = 0s and slightly reduced increase after t_m_ = 1-2s. Based on our model, only a small variation of free parameters fit to the experimental analysis, which assumes a linear adding up of molecular springs constants. Here, the first phase is determined by an increase in contact area within t_0_ = 1s and at a small maximal bead indentation of h_max_ = 80nm, corresponding to a maximal surface area of A_cap_ = 0.25µm. The second phase is determined by the low receptor diffusion of D_rec_ = 0.05 .. 0.07 µm^2^/s at a receptor density c_0_ = 4/µm^2^, which was also used for the BD simulations.

## DISCUSSION

In the following section, we discuss our experimental approach with the underlying concept and the results presented in this paper. Results include the analytical binding model, the Brownian dynamics (BD) simulations, the measured stepwise increase in binding strength and the averaged linear increase in binding strength.

### Experimental approach and concept

The usefulness of Photonic Force Microscopy (PFM) especially in combination with back focal plane (BFP) interferometric scattering has been demonstrated in many publications. Decisive advantages relative to alternative interferometric tracking techniques such as iSCAT [36] are a) that 3D tracking is well established, b) that the manipulation (trapping) and the tracking beam typically is the same [37], c) tracking at MHz rates as possible and that even in the presence of other scatterers 3D tracking is possible with nanometer precision [38].

Bead-based essays are commonly used to investigate interactions on a molecular scale especially between receptors and ligands. Based on this approach, only the technical advantages of PFM allow to reliably track the nanometers small changes in position fluctuations of optically trapped beads and to analyze them even in the presence of light-scattering cells. Although the presence of the cell influences the position signal at the QPD at low frequencies (typically < 10 Hz), the particle’s position signal fluctuations at > 10 kHz are not influenced by the cell [18]. This allows a reliable analysis of high-frequency fluctuations, which decay exponentially in most cases in their temporal autocorrelation, providing both the binding force constant (binding strength) and friction constant at subsequent time points of the experimental course. Because of the high sampling rate and 10^3^-10^6^ data points in time windows of 0.1 – 1 sec, changes in fluctuation widths of only Δx = 1 nm can be well distinguished.

However, at the current stage, orientation fluctuations of spherical particles cannot be measured with BFP interferometry [39, 40]. Although orientation fluctuations were estimated to be small, their influence on the measurement result could not be excluded, which required BD simulations.

### Brownian dynamics simulations

The performed BD simulation turned out to be very helpful, since they could answer a couple of questions, which could not be answered by the experiments. The first question is whether orientation fluctuations influence the center position tracking of the 1 µm large particle, when one or several receptors bind to the bead surface in a distance of *R*_*0*_ = 0.5 µm. It turned out that the hindered orientation fluctuations do not influence the strength of the position fluctuations, since the output values for the binding force constant are close to the input values of the simulation. Second, it could be shown by the simulations (Figure 3) that the position fluctuations of the membrane and thereby of the receptors hardly influence the binding force constant. However, it turns out that pure analyses of the fluctuation widths AC(τ=0) provide a noticeable deviation from the input values, while correct results are obtained by analysis of the full autocorrelation decay AC(τ). This answers a third question: a dominant reason for the superior τ method is the larger amount of particle positions considered by correlating them at different delay times. Furthermore, there is a separation of time scales since slow movements of the membrane occur typically on longer time scales than the short autocorrelation times of optical traps and molecular bonds. In the simulation results presented in Figure 4, we could demonstrate an increase of bound receptors with time, which consists of an initial indentation phase and a linearly increasing receptor diffusion phase and which coincides qualitatively and quantitatively with the experiments. In addition it strengthens the assumptions initial indentation and receptor diffusion - made for the analytical model according to eq. (4).

### Stepwise change in fluctuation width

The stepwise changes in fluctuation widths shown in Figure 4 and Figure 5 were obtained by two independent measurement series performed at different setups demonstrating the reproducibility of the results, which is not self-evident with this type of experiments. Based on the theoretical approaches described above, especially on linear restoring forces and quasi-thermal equilibria, the total force constants analyzed within time windows of 25 ms to 200 ms were quite similar with only 20% variation. Whereas changes in the position traces (recorded at 1 MHz) are hardly visible by pure eyes, the stepwise increase in stiffness (Figure 4A) and friction (Figure 4B) is clearly visible for the first two binding events. Here, the pure analysis of the fluctuation width (correlation amplitude at τ = 0, light blue curve) results in less pronounced steps than in the case of an analysis where time delays τ < 100 µs are considered (dark blue curve). This observation coincides very well with the Brownian dynamics simulations (autocorrelation-method, Figure 2C) and can be explained by the multifold better statistics when correlating different time points with each other. A similar behavior can be observed for the stepwise increase in the friction factor, which is better visible for larger time delays (τ < 100 µs, dark green curve) than for short time delay correlations in Figure 4B (τ < 30 µs, bright green curve). We interpret the increase in friction and damping by an increased viscous drag of the bead at the cell membrane or even the pericellular matrix (PCM).

The hypothesis that the change in fluctuations is not caused by single receptor bindings, but by sudden movements of the cell periphery such as PCM filaments or filopodia, cannot be fully excluded. While filopodia were not detected in the DIC images, the small PCM filaments cannot be resolved with conventional microscopy. However, contact with the soft PCM is likely to result in a smooth increase in stiffness and friction, but not in a jump like behavior [41].

Another hypothesis is that a bead movement out of the linear tracking regime results in a smaller fluctuation amplitude and thereby an erroneous increase in binding stiffness. Based on further BD simulations described in the supplementary material (Suppl. Figure S1), this effect was not relevant as long as the temporal decay of the AC(τ) was analyzed and not the pure fluctuation amplitude at τ = 0. The BD simulations also revealed (Suppl. Figure S2) that a smooth increase in viscosity and viscous drag of the bead towards the membrane [18] does not influence the correct analysis of the stiffness.

The exemplary experimental results of Figure 5 show a stepwise increase of the binding stiffness independent of two different smoothing strengths, similar to Figure 4B. Upon each binding, a linear decay of the stiffness constant (i.e. the measured autocorrelation time τ = γ/κ) over two seconds is visible, which may be explained by an increase of the friction constant γ, resulting from a soft embedding of the bead into the PCM. Distinct jumps in stiffness of κ_rec_ = 35 pN/µm and 50 pN/µm reflect the variety of the measurement results, and may appear larger than the average stiffness jump by effects such as the aforementioned decay in κ. The variety of binding strengths of totally 45 jumps in κ_tot_ is summarized by the histogram in Figure 5B, which reveals the most frequent stiffness jump at 19 pN/µm. Although measured higher stiffnesses can be a result of measurement uncertainty, the occurrence of stiffness jumps at a multiple of 19 pN/µm lead to the hypothesis that two or three binding events happen at the same time.

It will be of great interest for future experiments to correlate individual binding events measured through thermal position fluctuations with the underlying reorganization of the cytoskeleton using novel super-resolution microscopy methods operating at 100 Hz [42].

## CONCLUSION

We could observe for the first time single, successive binding events for IgG-coated beads to membrane receptors of living J774 macrophages using a label-free approach. The changes in the bead’s thermally driven position fluctuations during each binding event were tracked with quadrant photo diodes at rates of 10 kHz to 1 MHz. All experimental results coincide well with Brownian dynamics simulations and a simple analytical model, strengthening our interpretation of the results. We conclude that thermal noise encodes a variety of interactions that change with time especially in living systems. Artificial probes such as coated beads turn out to be once more well suitable to measure molecular interactions at their surfaces, since force transduction through the stiff bead and position tracking are very efficient. Future fluctuation binding experiments in combination with fluorescently labeled specific receptors might even reveal differences in the binding behavior under near physiological conditions, which may open new doors in toxicology, pharmacology and immunology.

## Supporting information

BrownianDynamics - Single Receptor - Binding Event

BrownianDynamics - Several receptors - slow approach

## ACKNOWLEDGMENT

We thank Dr. Felix Jünger and Dr. Rebecca Michiels for a careful reading of the manuscript and and helpful comments.

## AUTHOR CONTRIBUTIONS

AR and HK designed research; HK, TM and AR performed research; HK and TM analyzed data; AR and HK wrote the paper.

