## Supplementary Material for "Measuring stepwise binding of a thermally fluctuating particle to a cell membrane without labeling"

**To**

Alexander Rohrbach^1*^, Tim Meyer^1^, and Holger Kress^3*^

**1. Brownian dynamics simulations for nonlinear detector responses**

BD simulations allow to investigate the influence of the nonlinear detector response in QPD tracking caused by the finite with of the laser focus. The following figure illustrates the influence on the width of the position fluctuations, when the bead movement is in the nonlinear response regime of the detection system. The fluctuation width of an added stepwise bead movement will be underestimated, if the wrong analysis method is used.

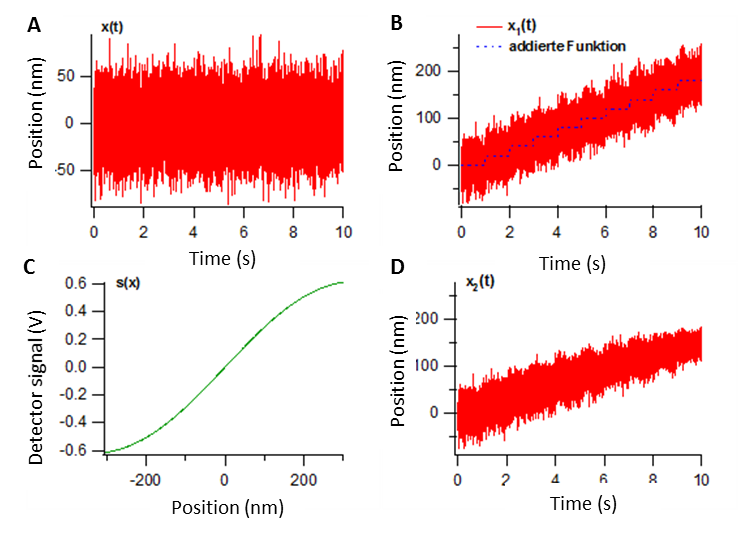

Figure S 1: BD simulations for nonlinear detector responses. (A) Particle trajectory and harmonic potential with ideal detector response and (B) particle trajectory with additional stepwise displacement. (C) Nonlinear detector response in QPD tracking. (D) Particle trajectory with additional stepwise displacement at nonlinear response.

**2. BD simulations with variable viscosity**

BD simulations can help to investigate the influence of the local increase of the bead’s viscous drag towards a cell membrane (Jünger, Kohler et al. 2015). The following Figure S2A shows that the corner frequency κ_trap_/γ_sol_ can be well reproduced by the BD simulations for a constant viscous drag γ_sol_ in the cell solution. However, for a increasing viscous drag γ_sol_(y) towards the cell, only the AC-time analysis method provides the correct results.

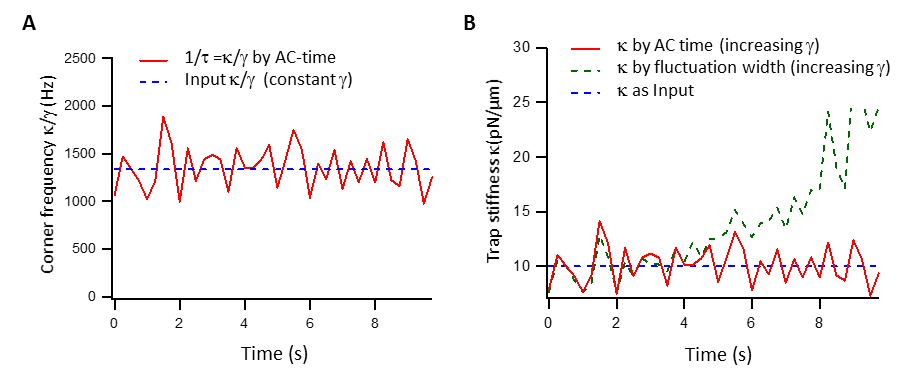

Figure S 2: BD simulation with nonlinear viscous drag close to a cell. The bead is approached to the cell in steps of 0.25s, corresponding to 40 steps over 10s. (A) The corner frequency, i.e. the ratio κ/γ can be well produced by AC analysis. (B) The stiffness κ can be reproduced by analyzing the autocorrelation time, but deviates from the input value if the fluctuation width σ = (kT/κ)^1/2^ is analyzed.

**3. Single binding BD simulations with variable viscosity**

The results of BD simulations demonstrate in the following figure that the increase of the viscous drag in the vicinity of the cell does not affect the jumps in position and stiffness (binding strength) from single receptor binding events.

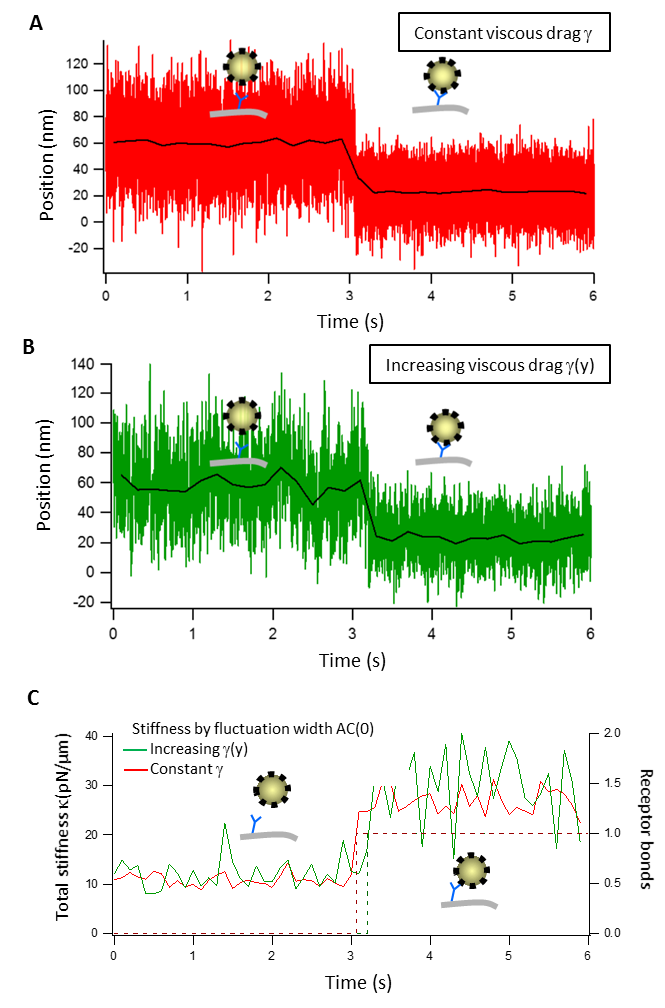

Figure S 3: Single binding BD simulation with nonlinear viscous drag at the membrane. The bead is approached to the cell in steps of 0.25s. (A) Position fluctuations before and after binding at constant viscous drag. The black line represents the 200ms average. (B) Position fluctuations before and after binding with increasing viscous drag. (C) Analysis of the total stiffness (receptor bonds) before and after binding for both behaviors of viscous drag. Dashed lines indicate the simulated number of receptors.
